# Integration of the Drug-Gene Interaction Database (DGIdb) with open crowdsource efforts

**DOI:** 10.1101/2020.09.18.301721

**Authors:** Sharon Freshour, Susanna Kiwala, Kelsy C. Cotto, Adam C. Coffman, Joshua F. McMichael, Jonathan Song, Malachi Griffith, Obi L. Griffith, Alex H. Wagner

**Affiliations:** Department of Medicine, Division of Oncology, Washington University School of Medicine, St Louis, MO, 63110, USA; McDonnell Genome Institute, Washington University School of Medicine, St Louis, MO, 63108, USA; Department of Genetics, Washington University School of Medicine, St Louis, MO, 63110, USA; Siteman Cancer Center, Washington University School of Medicine, St Louis, MO, 63110, USA; The Steve and Cindy Rasmussen Institute for Genomic Medicine, Nationwide Children’s Hospital, Columbus, OH, 43215, USA; Department of Pediatrics, The Ohio State University College of Medicine, Columbus, OH, 43210, USA

## Abstract

The Drug-Gene Interaction Database (DGIdb, www.dgidb.org) is a web resource that provides information on drug-gene interactions and druggable genes from various sources including publications, databases, and other web-based sources in one resource. These drug, gene, and interaction claims are normalized and grouped to identify aliases, merge concepts, and reduce redundancy. The information contained in this resource is available to users through a straightforward search interface, an application programming interface (API), and TSV data downloads. DGIdb 4.0 is the latest major update of this database. Seven new sources have been added, bringing the total number of sources included to 41. Of the previously aggregated sources, 15 have been updated. DGIdb 4.0 also includes improvements to the process of drug normalization and grouping of imported sources. Other notable updates include further development of automatic jobs for routine data updates, more sophisticated query scores for interaction search results, extensive manual curation of interaction source link outs, and the inclusion of interaction directionality. A major focus of this update was integration with crowd-sourced efforts, including leveraging the curation activities of Drug Target Commons, using Wikidata to facilitate term normalization, and integrating into NDEx for producing network representations.

## INTRODUCTION

Originally released in 2013, the Drug-Gene Interaction database (DGIdb)(1) serves as a single resource joining information on drug-gene interactions and druggability from multiple diverse sources. Previously, DGIdb has released two major updates, DGIdb 2.0(2) (in 2016) and 3.0(3) (in 2018) that included improvements to the user interface and search response times, the addition of an API, the introduction and improvement of gene and drug grouping methods, and the expansion of source content through the inclusion of new sources and updates of existing sources. Since the release of DGIdb 3.0, many of the existing sources have been substantially updated and new sources have become available. To help ensure that DGIdb offers current information, automatic updates have been further developed for multiple sources and a new background job management system (Sidekiq, sidekiq.org) has been implemented for routine job scheduling in the latest DGIdb release (DGIdb 4.0), described here. In addition to updates and integration of new sources, DGIdb 4.0 focuses on numerous improvements of search results, including an updated query score for interaction results and improved drug normalization. With these improvements, we have also made an effort to integrate crowd-sourced data and sources in several areas, including the addition of the crowd-sourced Drug Target Commons(4) as a drug-gene interaction source, and the use of the open, community Wikidata(5) resource for drug normalization. We also illustrate the value of our integration efforts in downstream community tools, through the incorporation of our data into NDEx(6).

### NEW AND UPDATED CONTENT

In an effort to ensure that DGIdb offers diverse and contemporary information, we have both updated many existing sources and added several new sources to DGIdb 4.0. For new sources, we have added two new sources of drug-gene interaction records and five new sources of druggable gene category records. The new sources of drug-gene interactions are Drug Target Commons(4) and COSMIC(7) (Supplemental Table 1). Drug Target Commons provides an extensive database of drug-gene interactions that have been curated through crowd-sourcing curation of expert knowledge. COSMIC provides an additional source of curated information on cancer mutations and drug resistance. The new category sources are Tempus xT(8), a panel of actionable cancer therapy target genes, a list of the top priority genes of the Illuminating the Druggable Genome (IDG)(9) Initiative, the Human Protein Atlas(10), Oncomine(11), a clinical cancer biomarker assay, and Pharos(12), which hosts data as part of the IDG program and focuses on understudied targets in three broad classes of druggable genes: G protein-coupled receptors, ion channels, and kinases (Supplemental Table 1). From these new sources, we have added 23,916 new drug-gene interaction records and 8,463 new druggable gene category records (Figure 2).

Additionally, we have also updated multiple sources including large, well-curated sources such as DrugBank(13), Guide to Pharmacology(14) Interactions, Gene Ontology(15), (16), OncoKB(17), PharmGKB(18), and the Therapeutic Target Database(19) (Figure 2, Supplementary Table 1). To facilitate routine updates, the importer for PharmGKB has been updated to an online importer that can be run periodically using DGIdb’s new automated job scheduling system (see Technology Improvements). Similarly, Pharos(12), one of our new sources, has been implemented as an online updater. Of the 41 sources in DGIdb, 12 sources are now imported using the online updater format, including Entrez(20), the core source of gene concepts for gene grouping in DGIdb, and Ensembl(21), a key source of gene aliases. In DGIdb 4.0, the number of genes imported from Entrez has increased from 41,102 to 42,767 and Ensembl has been updated from version 90_38 to version 101_38. We have also migrated several sources from domain-specific language (DSL) importers to the TSV importer style implemented in DGIdb 3.0.

In DGIdb 4.0, the database structure and presentation model was updated to allow sources to be imported with multiple source types. This change enables merging of sources that were previously duplicated for each independent claim type (drug, gene, interaction, druggable gene category). For example, we currently import both interaction claims and druggable gene category claims from Guide to Pharmacology(14) with two separate importers. Instead of listing two independent sources (GuideToPharmacologyInteractions, GuideToPharmacologyGenes), we can now have a single source (GuideToPharmacology). This is intended to simplify and unify claim sources to aid in downstream interpretation. Additionally, supporting sources that have multiple source types enables easy extension to collect more informative claim type information. For example, some claims from CIViC(22) can be imported with the additional category of drug resistance. Overall, these changes will increase the efficiency and accuracy of the process of importing and updating sources in DGIdb 4.0 compared to previous versions and will make it easier for users to evaluate individual sources.

### IMPROVED TRANSPARANCY AND DETAILS ON LICENSING OF SOURCES

In DGIdb 4.0, we have made a significant effort to update and improve the information we provide on licensing of sources imported into our database through manual curation of licensing types and details for every source. This information is now readily available on the sources page and sources with special licenses are noted on the downloads page (Figure 1). Since DGIdb 3.0, several existing sources have made changes to their licensing, making data from some sources more broadly available and data from other sources more restricted. Notably, PharmGKB(18) has moved to a more permissive Creative Commons Attribution-ShareAlike 4.0 International License and DrugBank(13) has adopted a custom non-commercial license. In contrast, OncoKB(17) has restricted API access to registered/approved non-commercial research use only, and JAX-CKB(23) has restricted API access to negotiated licenses only. Both resources continue to provide access to a portion of their data for free through their respective web clients. When exporting data TSVs for downloading, we partition data with restricted license types separately and note the details of licensing. If data cannot be made available for downloading, users are given directions on how data can be obtained from the sources themselves.

**Figure 1:**
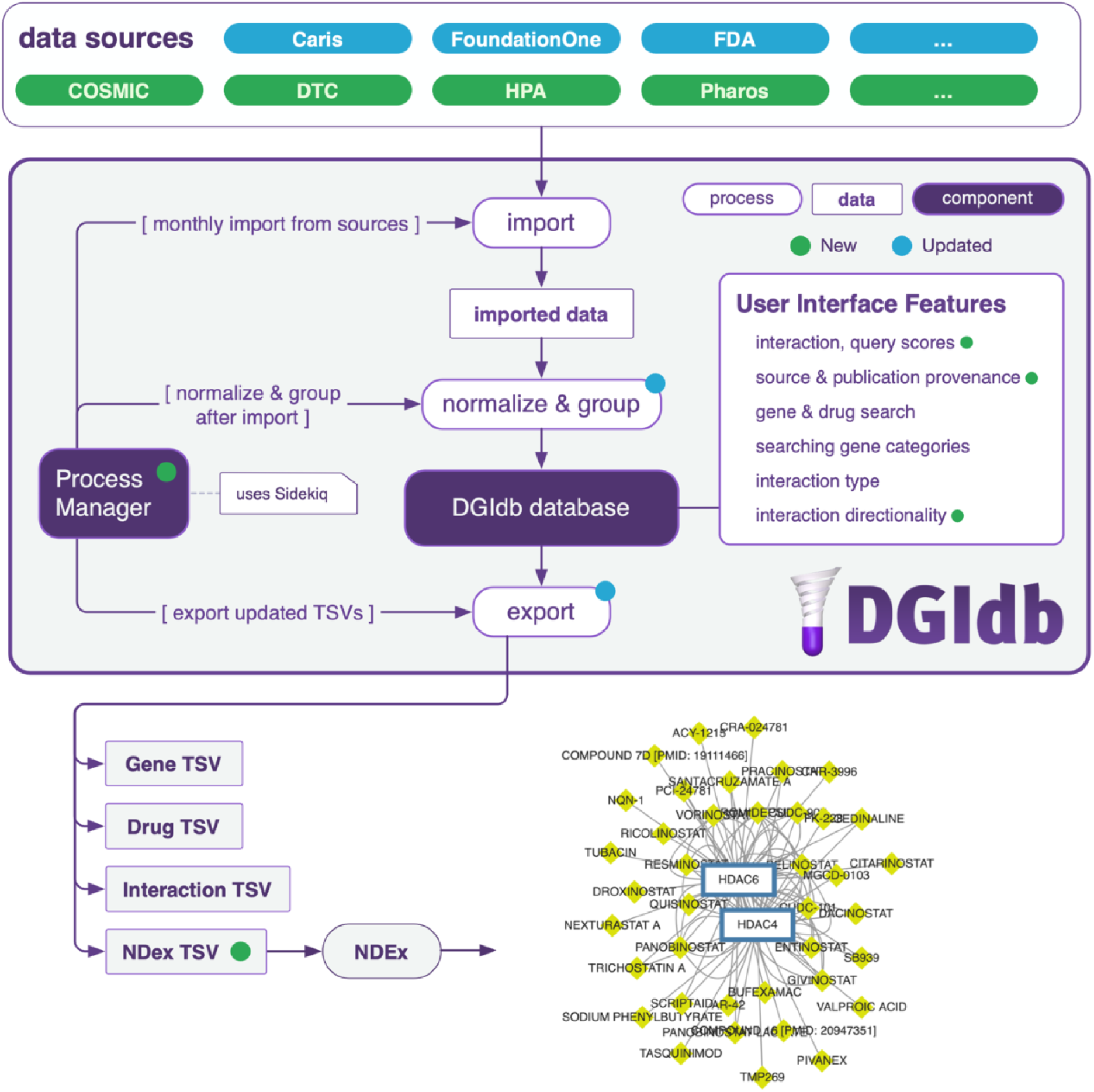
Overview of main components of DGIdb. Data sources are imported from outside resources (over 40 as of DGIdb 4.0), normalized and grouped with internal processes to prepare records to be displayed in DGIdb, and exported to TSV downloads or integrated with other resources. Process management is handled by Sidekiq for automation of importing, normalization and grouping, and exporting. A subset of new data sources are highlighted in green, a subset of updated pre-existing data sources are highlighted in blue. The updated sources highlighted in this figure are some of the sources that have been updated through manual curation. Information on additional sources and their status in DGIdb 4.0 can be found in Figure 2 and Supplemental Table 1. New features and technologies from DGIdb 4.0 are indicated with green dots, pre-existing features and technologies that have been updated are indicated with blue dots.

**Figure 2:**
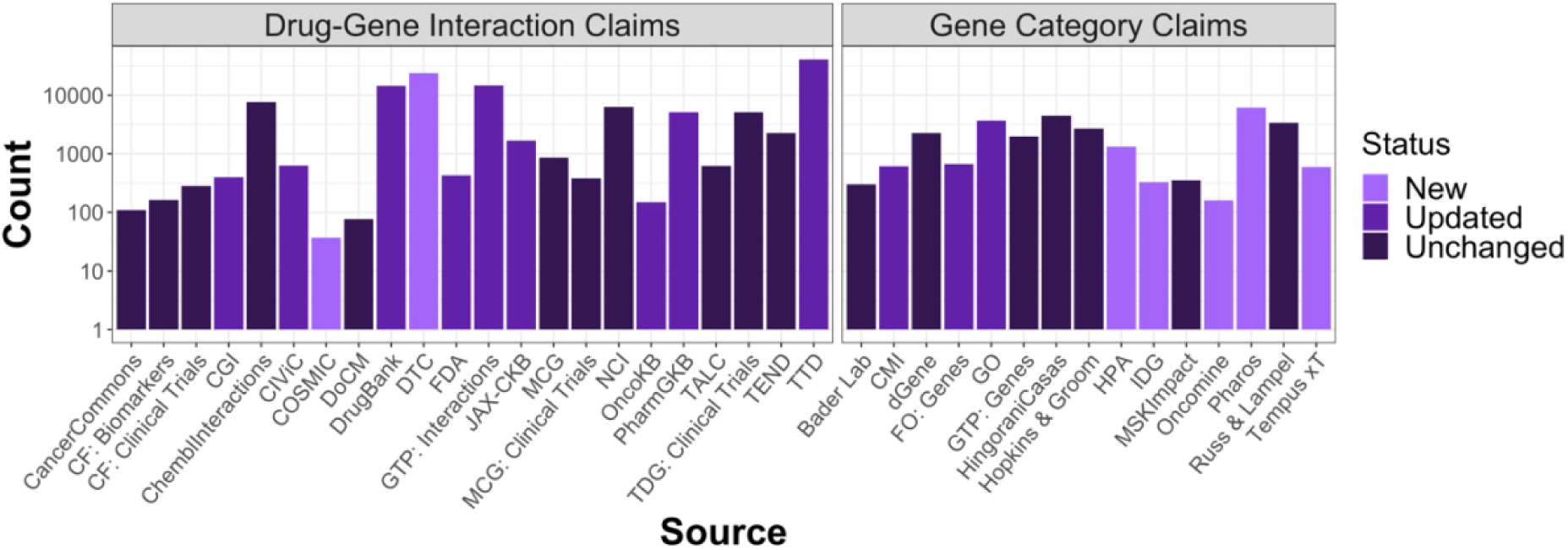
DGIdb 4.0 content by source. The number of drug-gene interaction claims (first panel) and druggable gene categories (second panel) are separated into three categories: sources that are new, sources that existed in the DGIdb previously but have been updated for 4.0, or sources that existed previously but have not been updated. Abbreviations: CF = Clearity Foundation, CGI = Cancer Genome Interpreter, CMI = Caris Molecular Intelligence, FO = Foundation One, GO = Gene Ontology, GTP = Guide to Pharmacology, HPA = Human Protein Atlas, IDG = Illuminating the Druggable Genome, JAX-CKB = JAX-Clinical Knowledgebase, MCG = My Cancer Genome, MSK = Memorial Sloan Kettering, OncoKB = Precision Oncology Knowledge Base, TALC = Targeted Agents in Lung Cancer, TDG = The Druggable Genome, TEND = Trends in the Exploration of Novel Drug targets, and TTD = Therapeutic Target Database.

### MONTHLY DATA RELEASES

DGIdb 3.0 implemented online updaters that imported data from dynamic sources (such as CIViC(22), Guide to Pharmacology(14), OncoKB(17), etc.) periodically, usually monthly. As a result, the static TSV data releases available on our Downloads page would quickly become outdated. For DGIdb 4.0, we have implemented monthly data releases of these TSVs to coincide with monthly runs of the online updaters, to ensure that TSVs available for download reflect the most up to date information in our database (Figure 1). The Downloads page now makes available the current Gene, Drug, Interaction, and Category TSVs as well as historic TSVs from past runs. These serve as de-facto snapshots of the data in DGIdb over time.

### INTEGRATION WITH NDEX AND OTHER EXTERNAL RESOURCES

The Network Data Exchange (NDEx)(6) is a community resource that allows sharing and publishing of biological data in a network-based format. For DGIdb, integration with the NDEx platform provides a resource for the visual representation of relationships and interactions between drugs and genes present in our database, allowing users to easily explore a global overview of drugs and genes of interest. NDEx TSVs are generated monthly and automatically uploaded to the NDEx server to keep the DGIdb network in NDEx up-to-date (Figure 1). Previously, DGIdb has been integrated with other community resources such as GeneCards(24), which uses DGIdb as a source for interactions between GeneCards gene entries and drugs, and BioGPS(25), which offers a DGIdb plug-in for searching drug-gene interactions. Since the publication of DGIdb 3.0, DGIdb has been regularly integrated with additional resources such as CancerTracer(26), Gene4Denovo(27), SL-BioDP(28), TargetDB(29), and OncoGemini(30).

### UPDATED QUERY AND INTERACTION SCORES

One of the main features added in DGIdb 4.0 is the concept of relative query scores for interaction search results. Previously, interaction search results displayed only a static score based on evidence of an interaction (i.e. the number of publications and sources supporting an interaction claim). This score did not take into account whether the gene and drug involved in a given interaction were also part of a large number of other interactions and, thus, had a low specificity that should be penalized. In addition, when searching for a set of genes or drugs, the previous interaction score would also not prioritize results with overlapping interacting drugs or genes, which might be of more interest to the user, particularly in drug discovery and pathway applications.

DGIdb 4.0 now provides a query score that is relative to the search set and considers the overlap of interactions in the result set. For interaction searches using a gene list, the query score depends on the evidence scores (publications and sources), the number of genes from the search set that interact with the given drug, and the ratio of average known gene partners for all drugs to the known partners for the given drug (Figure 3). Similarly, for interaction searches using a drug list, the query score depends on the evidence scores (publications and sources), the number of drugs from the search set that interact with the given gene, and the ratio of average known drug partners for all genes to the known partners for the given gene (Figure 3). In effect, this means that genes and drugs with many overlapping interactions in the search set will rank more highly, with the caveat that drugs or genes involved in many interactions, in general, will have lowered scores (Figure 3).

**Figure 3:**
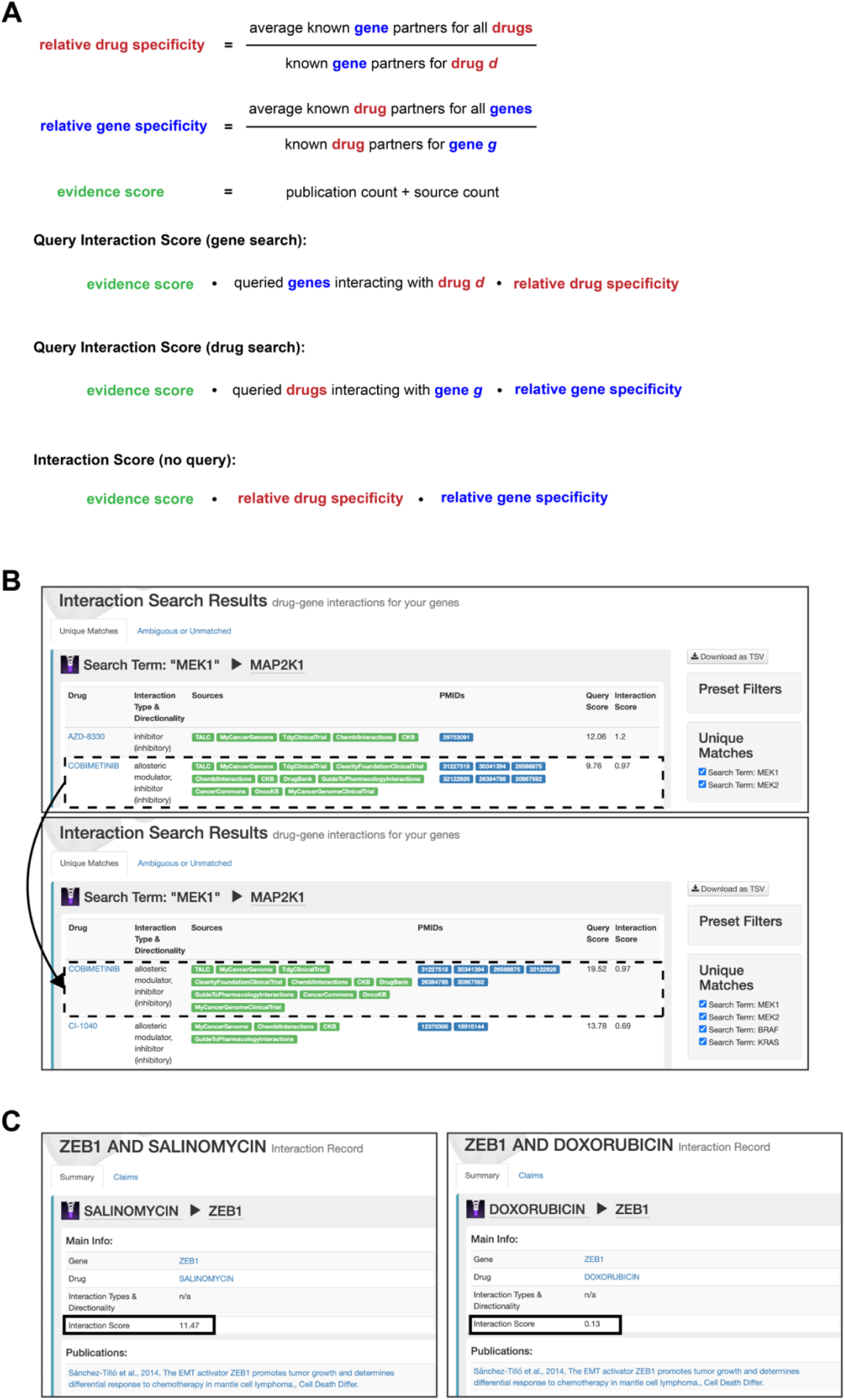
Overview of DGIdb’s new query interaction scores and interaction scores. **A)** Schematic of how each of the new scores is calculated within DGIdb. Gene and drug queries both return a query interaction score that is dependent on the search terms. Each interaction has a score that is calculated independently of other search terms. **B)** Example of a query interaction score changing based on the terms searched. In the first panel, only *MEK1* and *MEK2* were searched and the score for the interaction between *MEK1* and Cobimetinib was 9.76. In the second panel, *BRAF* and *KRAS* were added to the search query. These both interact with Cobimetinib and thus raise the query interaction score. **C)** Example of interaction score. The panel on the left shows the interaction between *ZEB1* and Salinomycin. This is only interaction for Salinomycin and thus it has a high interaction score. The panel on the right shows the interaction between *ZEB1* and Doxorubicin. Doxorubicin is involved in 87 interactions within DGIdb and thus has a much lower interaction score.

The static interaction score has also been improved. It now mirrors the query score except that it removes the dependency on queried gene or drug sets and only uses evidence scores, the ratio of average known gene partners for all drugs to the known partners for the given drug, and the ratio of average known drug partners for all genes to the known partners for the given gene (Figure 3). In summary, the introduction of the relative query score will provide users with a score that gives a more intuitive ranking of drugs or genes based on the search set of interest, allowing the prioritization of drugs or genes that have overlapping, specific interactions with the search set. Similarly, the improvements to the static interaction provide a more nuanced scoring system that takes into consideration the number of interactions for a drug-gene pair, in addition to the previous evidence scores, giving a more informative static score.

### OTHER SEARCH RESULT IMPROVEMENTS

Another major focus of the DGIdb 4.0 update is the improvement of features related to search results and information available on the user interface (Figure 1). One such improvement is the addition of link outs to specific interaction evidence, where available. These will allow users to browse to a specific source page for an interaction claim which might provide more information than available on DGIdb.

We have also added information on the directionality of interaction types. This directionality indicates whether the interaction type of a specific interaction result has an inhibitory or activating effect overall (when the directionality is available for an interaction type). While the directionality may be clear for some interaction types, e.g. activators have activating directionality, some interaction types are less intuitive, e.g. chaperones have activating directionality. Inclusion of directionality can make it easier for users to distinguish interactions that are more relevant for their purposes. For example, a user interested in exploring drugs that inhibit a particular gene will readily see that chaperone interactions are activating, without further investigation of the interaction, and might ignore those records.

Finally, while we introduced drug and gene filters to the interaction search view in the last major update, we have had several requests to define how these filters are implemented. To address this lack of transparency on the UI, we have now added a link to the FAQ page where these filters are now defined.

### IMPROVEMENT OF DRUG NORMALIZATION APPROACH

In addition to the improvement of features related to search results, another main focus of the DGIdb 4.0 update is the improvement of drug grouping and normalization. Previously, we grouped drug claims using a rule-based pairwise association approach. This process was cumbersome, requiring a lengthy and complete re-grouping of all claims whenever we updated sources in order to generate consistent groupings. We revised this approach by creating a normalization component independent of the claims aggregated by DGIdb, that could be run on a per-source basis. When redesigning this part of DGIdb, we took steps to enable reuse of this normalizer as a modular component for other resources. To this end, we leveraged and contributed to a drug normalization service from the Variant Interpretation for Cancer Consortium (VICC) (the “*thera-py*” Python Package; source code online at https://github.com/cancervariants/therapy-normalization). Among our contributions to *thera-py* was a normalizer for the Wikidata(5) resource, further enabling community contributions to assist in concept normalization both for DGIdb and other resources reliant upon the VICC normalization services.

Drug claims from DGIdb were normalized using the ChEMBL and Wikidata normalizers from *thera-py*. Rules were written to formalize grouper behavior based upon match characteristics of a query. Briefly, these rules prioritize matches to primary labels over aliases, exact case over case-insensitive, and ChEMBL over other normalizers. An algorithm for constructing a merged drug concept from normalizer results was specified, enabling a standardized set of aliases for a given concept identifier. Pseudocode for this algorithm is provided (see Supplementary Data), and all implemented code is available on our public repository (see Data Availability).

### TECHNOLOGY IMPROVEMENTS AND IMPLEMENTATION

To handle increased web traffic and integration with other tools, we have upgraded DGIdb to Rails 6 (from Rails 5), upgraded to Ruby 2.6.5 (from Ruby 2.3), upgraded to PostgreSQL 12 (from PostgreSQL 9.6), and upgraded the server to the latest Ubuntu LTS release (20.04). In addition to the new features and performance benefits these upgrades bring, they will ensure that we continue to remain on supported software versions that receive regular security updates.

In order to keep DGIdb’s underlying source data current, we had previously implemented an automated job scheduling framework using DelayedJob to schedule monthly runs of online updaters. In this release, we switched to using Sidekiq (Figure 1). In contrast to DelayedJob, Sidekiq offers a convenient user interface which makes identifying job failure reasons and rescheduling of failed jobs easier. Furthermore, the addition of Airbrake, an online tool for exception tracking, gives error reviews and notifies the development team of these errors in real-time (for instance, via email).

To ease future implementation of fixes and new features, we moved testing to a GitHub continuous integration (CI) workflow which allows us to continuously test newly committed code for errors against multiple versions of Ruby and PostgreSQL.

### SUMMARY AND FUTURE DIRECTIONS

With our most recent release, DGIdb has received significant improvements to source content, functionality such as searching and grouping, and underlying application technology. We have significantly expanded the number of records in our database through the addition of new sources and updates of existing sources. Furthermore, we have improved our ability to maintain regular content updates through the implementation of additional online importers for several sources and the use of Sidekiq for automatic job processing. We have revised our process for drug grouping and normalization to be batched by resource and to leverage continual improvement through community contributions to the VICC *thera-py* normalizers and the Wikidata public-domain crowdsourcing platform. Finally, several important changes have been made to search results and the information presented on the UI. We have implemented more sophisticated notations of relative and static interaction scores, improved the relevance of interaction source link outs wherever possible, and included the concept of directionality for interaction results.

Although the updates in DGIdb 4.0 have improved the usability and content of our resource, we expect there will still be a need for future improvements. One technology improvement on our roadmap is converging the public facing API with the internal code that powers the web views. Ultimately, we want the APIs available for general use to be the same ones powering our HTML pages. This would provide an even more fully featured API to end users while reducing our overall maintenance burden by eliminating redundant code. Additionally, one possible feature to add to the user-interface is displaying information on related genes on the gene page for a given gene. Finally, we also plan to continue updating sources to online updaters, where possible, and migrating TSV-based sources from the legacy DSL importer style to the updated DGIdb 3.0 TSV importer style.

## AVAILABILITY

DGIdb is an open access database and web interface (www.dgidb.org) with open source code available on GitHub (https://github.com/griffithlab/dgi-db) under the MIT license. We also provide data downloads for drug claims, gene claims, and interaction claims on the website in addition to a SQL data dump (http://dgidb.org/downloads). Information about the API and its endpoints can also be found on the website (http://dgidb.org/api).

## SUPPLEMENTARY DATA

Supplementary Data are available at NAR online.

## ACKNOWLEDGEMENTS

We want to thank the creators and maintainers of the seven new resources added to DGIdb and the many previously incorporated resources, as well as our growing community of users for notifying us of minor and major issues and for their suggestions on new features and improvements to DGIdb. We would also like to thank the members of the NDEX team, and in particular Dexter Pratt and Rudolf Pillich for their efforts in integrating DGIdb data into the NDEX resource.

## FUNDING

This work was supported by the National Human Genome Research Institute [K99HG010157 to A.H.W. and A.C.C., R00HG007940 to M.G.] and the National Cancer Institute [U01CA209936, U01CA248235, U24CA237719 to M.G., O.L.G., S.K., A.C.C., A.H.W. and J.F.M.]. Funding for open access charge: departmental funding

## CONFLICT OF INTEREST

None declared

## Notes

### Competing Interest Statement

The authors have declared no competing interest.

