## Supplementary Data for "Integration of the Drug-Gene Interaction Database (DGIdb) with open crowdsource efforts"

### Supplemental Table 1: Description of Sources in DGIdb 4.0

| Data Type | Source | Description of Sources and Imported Data | 4.0 Status |
|---|---|---|---|
| Drug Gene Interactions | CancerCommons(Shrager, 2011) | CancerCommons provides a number of drugs that are approved or undergoing clinical trials for use in lung, prostate, and skin cancer. Drug-gene interactions were extracted from their web pages describing these diseases and imported. | P |
| Drug Gene Interactions | Cancer Genome Interpreter (CGI) (https://www.cancergenomeinterpreter.org/) | CGI is a database that provides information about identified alterations and currently available treatment treatment. The biomarkers per variant file was downloaded and parsed for import into the DGIdb. | A, U |
| Drug Gene Interactions | ChEMBL: Interactions(Bento, 2014) | ChEMBL is a database of small molecules capable of bioactivity that have been annotated with metadata such as 2-D structures and calculated properties (e.g. Molecular Weight). Drug-gene interactions were pulled from the database. | P |
| Drug Gene Interactions | CIViC(Griffith, 2017) | CIViC is a community-driven platform for identifying actionable variants in cancer. CIViC genes were pulled via the provided API and drug-gene interactions of those genes were imported. | A, U |
| Drug Gene Interactions | Clearity Foundation: Biomarkers | The Clearity Foundation has analyzed thousands of tumors from ovarian cancers and retrospectively identified biomarkers that predict treatment response to select drugs in patient ovarian tumors. These interaction data were extracted from their web page and imported. | P, S |
| Drug Gene Interactions | Clearity Foundation: Clinical Trials | 124 curated clinical trial records were provided by The Clearity Foundation based on their relevance to breast and ovarian cancer. The interactions from these trials were imported. | P, S |
| Drug Gene Interactions | COSMIC(Tate, 2019) | Cosmic provides curated information on cancer mutations and drug resistance. A CSV file was manually created from the table available at https://cancer.sanger.ac.uk/cosmic/drug_resistance for import in DGIdb. | N |
| Drug Gene Interactions | DoCM(Ainscough, 2016) | DoCM is a manually curated database of mutations associated with cancer progression. Drug-gene interaction data were pulled via the API and imported into the DGIdb. | A, P |
| Drug Gene Interactions | DrugBank(Wishart, 2018) | DrugBank is a large community resource detailing drug to drug target information. The full DrugBank database was downloaded as an xml file and parsed for import into the DGIdb. | A, U |
| Drug Gene Interactions | Drug Target Commons(Tang, 2018) | DTC is an extensive database of drug-gene interactions that have been curated through crowd-sourcing curation of expert knowledge. This data was downloaded from the Drug Target Commons website as a raw text preprocessed to only keep entries that have a compound_name, gene_names, and have a value of "active" or "Active" in the activity_comment column before import into DGIdb | N |
| Drug Gene Interactions | FDA Biomarkers (https://www.fda.gov/drugs/science-and-research-drugs/table-pharmacogenomic-biomarkers-drug-labeling/) | The FDA provides drug-gene interactions in their Pharmacogenomic Biomarkers in the Drug Labeling table. 427 drug-gene interactions were pulled from this table for various diseases. | U |
| Drug Gene Interactions | Guide To Pharmacology: Interactions(Southan, 2016) | The Guide to Pharmacology is an extensive resource detailing pharmacological targets and their corresponding drugs. The online updater downloads the interactions.tsv file and then parses it for import into the DGIdb. | A, U |

| | | | |
|---|---|---|---|
| Drug Gene Interactions | Jax-Clinical Knowledgebase (JAX-CKB, Patterson, 2016) | JAX-CKB is a knowledge base that provides drug-gene interactions based on a tumor's genomic profile as well as drug efficacy and resistance evidence. Drug-gene interactions and metadata were pulled via the API and imported into the DGIdb. | A, U |
| Drug Gene Interactions | My Cancer Genome(Yeh, 2013) | My Cancer Genome provides information on how mutations drive cancer, and the implication of those mutations for treatment, including linking interactions between specific mutations and therapies. These interactions were extracted from their website with permission. | P, S |
| Drug Gene Interactions | My Cancer Genome: Clinical Trials(Yeh, 2013) | My Cancer Genome previously supported the searching of clinical trials to support their interactions. When this was available, these data were extracted from their website with permission and imported into the DGIdb. | P, S |
| Drug Gene Interactions | National Cancer Institute (NCI) Cancer Gene Index (https://wiki.nci.nih.gov/display/cageneindex) | This resource was curated through scraping interactions from publications followed by manual curation. The Gene-Compound xml file was downloaded and parsed for inclusion into the DGIdb. | P, S |
| Drug Gene Interactions | Precision Oncology Knowledge Base (OncoKB)(Chakravarty, 2017) | OncoKB is a knowledgebase that provides information about the specific effects of a somatic molecular alteration and potential treatment options with predicted drug response information. Drug-gene interactions and metadata were pulled via the API and imported into the DGIdb. | A, U |
| Drug Gene Interactions | PharmGKB(McDonagh, 2011) | PharmGKB is a knowledge resource that collects information about potentially clinically actionable gene-drug associations. These associations were downloaded via the relationships.tsv file from PharmGKB for parsing and import into the DGIdb. | A, U |
| Drug Gene Interactions | Targeted Agents in Lung Cancer (TALC)(Somaiah, 2014) | This 2012 publication presented a comprehensive survey of targeted agents in lung cancers. The data were provided as pdf tables within the publication and were manually reviewed for import into the DGIdb. | P, S |
| Drug Gene Interactions | TDG Clinical Trials(Rask-Andersen, 2014) | This 2014 publication evaluated drug-target interactions in the CenterWatch Drugs in Clinical Trials Database. The aggregated drug-gene interactions were provided as a supplementary table, which was manually reviewed for import into the DGIdb. | P, S |
| Drug Gene Interactions | Trends in the Exploitation of Novel Drug Targets (TEND)(Rask-Andersen, 2011) | This 2011 publication was an extensive manual curation of FDA approved drugs and their targets from DrugBank. The results of this effort are in a supplementary table, which was manually reviewed for import into the DGIdb. | P, S |
| Drug Gene Interactions | Therapeutic Target Database (TTD)(Zhu, 2012) | The periodically updated TTD resource provides information about therapeutic targets and their corresponding drugs. This information was downloaded from the TTD website as raw text, and then parsed for import into the DGIdb. Drugs that were listed as 'Terminated', 'Withdrawn from market', or 'Discontinued' were not included. | U |
| Drug Definitions | ChEMBL:Molecules(Bento, 2014) | ChEMBL is a database of small molecules capable of bioactivity that have been annotated with metadata such as 2-D structures and calculated properties (e.g. Molecular Weight). ChEMBL serves as a source of drug concepts for normalization. | A, U |
| Drug Definitions | PubChem(Kim, 2016) | PubChem served as a canonical concept for drugs in the DGIdb 3.0. It has been removed in 4.0 as the updated drug grouper relies on normalized concepts from ChEMBL and Wikidata | R |
| Drug Definitions | Wikidata(Vrandečić, 2014) | Wikidata is a free and open knowledge base containing structured data that can be read and edited by both humans and machines. Wikidata serves as a source of drug concepts for normalization. | A, N |
| Gene Definitions | Ensembl(Yates, 2016) | Ensembl gene IDs were imported and linked to Entrez gene records. These were imported from Ensembl's ftp site using the transcript GTF file, | A, U |

| | | | |
|---|---|---|---|
| | | and also the NCBI gene_info file. | |
| Gene Definitions | NCBI Entrez Gene(Brown, 2015) | Entrez gene serves as the canonical concept for genes in the DGIdb. Entrez Gene records are imported from NCBI using the online gene_info file. | A, U |
| Druggable Gene Categories | Bader Lab(Edwards, 2011) | A publication detailing four large protein families. These gene category claims are available from the supplementary data hosted online, which were downloaded for import into the DGIdb. | P, S |
| Druggable Gene Categories | Caris Molecular Intelligence(https://www.carismolecularintelligence.com/) | We extracted clinically actionable gene lists from the biomarker and NGS panels provided by Caris Life Sciences online at http://www.carismolecularintelligence.com/. | U |
| Druggable Gene Categories | dGENE(Kumar, 2013) | An annotation tool for checking if genes are members of one of ten druggable gene families. The lists of these gene families were provided by the authors and imported into the DGIdb. | P, S |
| Druggable Gene Categories | Foundation One Genes(Wagle, 2012) | Foundation One diagnostic test focuses on clinically actionable genes, which are available via an online table on their website. These data were extracted from the site for import into the DGIdb. | U |
| Druggable Gene Categories | GO(Gene Ontology, 2015) | Multiple gene categories were expertly selected and an online updater was created which queries the Gene Ontology API for these categories for import into the DGIdb. | A, U |
| Druggable Gene Categories | Guide To Pharmacology: Genes(Southan, 2016) | The Guide to Pharmacology is an extensive resource detailing pharmacological targets and their corresponding targets. The interaction and target and family files were downloaded and parsed for import into the DGIdb. | P |
| Druggable Gene Categories | HingoraniCasas(Finan, 2017) | A publication from the Casas lab that used computational approaches to identify druggable genes from genome-wide association studies. Interaction claims were curated from this paper's supplementary information. | P, S |
| Druggable Gene Categories | Hopkins & Groom(Hopkins, 2002) | Considered the 'original' druggable genome by the DGIdb 1.0, information on genes that were predicted to make good drug targets were extracted from the paper for import into the DGIdb. | P, S |
| Druggable Gene Categories | Human Protein Atlas(Uhlen, 2017) | Used a computer-based modeling approach to identify genes predictive of cancer outcome. A TSV was created by going to the https://www.proteinatlas.org/search/protein_class%3APotential+drug+targets page and selecting the columns Gene, Gene synonym, Ensembl gene id, Gene description, Uniprot accession, and Protein class (from the Gene section) and clicking "Download TSV". This TSV was then imported into DGIdb. | N |
| Druggable Gene Categories | Illuminating the Druggable Genome (IDG) Initiative(Rodgers, 2018) | IDG focuses on understudied targets in three broad classes of druggable genes: G protein-coupled receptors, ion channels, and kinases. A TSV file was created manually from the table available at https://druggablegenome.net/IDGProteinList for import into DGIdb. | N |
| Druggable Gene Categories | MSK IMPACT(Cheng, 2015) | Clinically actionable genes were extracted from the supplementary tables of the Memorial Sloan Kettering IMPACT paper. | P, S |
| Druggable Gene Categories | Oncomine(Williams, 2018) | A clinical cancer biomarker assay. A CSV was manually created from the table available in the Comprehensive Assay V3 flyer (https://assets.thermofisher.com/TFS-Assets/LSG/brochures/oncomine-comprehensive-assay-v3-flyer.pdf) for import into DGIdb. | N |
| Druggable Gene | Pharos(Nguyen, 2017) | Hosts data as part of the IDG program. An online updater was created, which queries the Pharos GraphQL API | A, N |

| | | | |
|---|---|---|---|
| Categories | | (https://pharos-api.ncats.io/graphql) for gene category data. | |
| Druggable Gene Categories | Russ & Lampel(Russ, 2005) | Considered the 'updated' druggable genome by the DGIdb 1.0, information on genes that were predicted to make good drug targets were sent by the authors for import into the DGIdb. | P, S |
| Druggable Gene Categories | Tempus xT(Beaubier, 2019) | Panel of actionable cancer therapy target genes. A TSV was manually created from the gene list published by Tempus (https://www.tempus.com/wp-content/uploads/2018/12/xT-Gene-List_112818.pdf) for import into DGIdb. | N |

A: Denotes source that is automatically updated, N: Denotes source that is new in the DGIdb 4.0,
P: Denotes source that was previously in the DGIdb 3.0 and has not been updated, R: Denotes a source that was previously in DGIdb 3.0 but has been removed in 4.0, S: Denotes source that is static (e.g. a table from a paper, a data download from a paper, or a source that has been deprecated), U: Denotes source that has been updated in the DGIdb 4.0.

### Pseudocode for the DGIdb drug grouper

***main***

1. Look for ChEMBL normalizer with match >= 80
    a. If match has only one record:
        i. return record normalized to chembl_id
    b. If match has multiple records all with same chembl ID:
        i. return record normalized to chembl_id
    c. If match has multiple records with different chembl IDs:
        i. select record with highest max phase
            1. if multiple records, narrow down to those with a trade name
                a. if still multiple records, use the one with the lowest chembl id and normalized to chembl_id
                b. if one record, normalized to chembl_id
2. Select normalizer(s) with highest match and match > 0:
    a. Further select on normalizers have records with chembl ids, if any
        i. if no normalizers have records with chembl ids, keep all normalizers
    b. Select results from normalizer with highest priority
        i. If selected normalizer is ChEMBL, go to Step 1a
        ii. If match has only one record:
            1. If match has a chembl id and id is valid
                a. return record normalized to chembl ID
            2. Else
                a. return record
        iii. if match has multiple records:
            1. select any records with a chembl id that is valid
                a. if the chembl ids are the same, return record normalized to chembl_id
                b. if the chembl ids are not the same:
                    i. normalize each record to chembl ID
                    ii. if only one record has highest max_phase:
                        1. return normalized record with greatest max_phase
                    iii. else:
                        1. if match score >= 40:
                            a. narrow down to those with a trade name
                                i. if still multiple records, use the one with the lowest chembl id and normalized to chembl_id
                                ii. if one record, normalized to chembl_id

            2. if match score < 40:
                a. do not normalize
        2. if no records have a chembl id
            a. do not normalize
3. If no normalizer(s) with match > 0:
    a. do not normalize

***records with chembl ids***

a record has a chembl ID as the concept_identifier from the chembl normalizer, or may contain one as a CURIE in the other_identifiers field namespaced with "chembl:"

***normalize record to chembl ID***

NOTE: it may be beneficial to cache this operation on a given chembl ID, as the result will not change.

1. run normalizer query on chembl ID
2. select all normalizers with match >= 80
    a. if a chembl record exists:
        i. create copy of chembl record to *new_record*
        ii. add *aliases* and *other_identifiers* from other normalizers to *new_record*
        iii. return *new_record*
    b. if a chembl record doesn't exist
        i. return None

***priority ranking***

1. ChEMBL normalizer
2. Wikidata normalizer
